# Visualization of Loki- and Heimdallarchaeia (Asgardarchaeota) by fluorescence *in situ* hybridization and catalyzed reporter deposition (CARD-FISH)

**DOI:** 10.1101/580431

**Authors:** Michaela M. Salcher, Adrian-Ştefan Andrei, Paul-Adrian Bulzu, Zsolt G. Keresztes, Horia L. Banciu, Rohit Ghai

## Abstract

Metagenome-assembled genomes (MAGs) of Asgardarchaeota are starting to be recovered from a variety of habitats, broadening their environmental distribution and providing access to the genetic makeup of this archaeal lineage. Despite their singular phylogenetic position at the base of the eukaryotic tree of life, the morphology of these bewildering organisms remains a mystery. In order to visualize this elusive group, we applied a combination of CARD-FISH and epifluorescence microscopy on coastal hypersaline sediment samples, using specifically designed CARD-FISH probes for Heimdallarchaeia and Lokiarchaeia lineages and provide the first visual evidence for both these groups. Here, we show that while Heimdallarchaeia are characterized by a uniform cellular morphology typified by central DNA localization, Lokiarchaeia display a plethora of shapes and sizes that likely reflect their broad phylogenetic diversity and ecological distribution.

## Main text

The discovery of the Asgardarchaeota not only revealed the closest archaeal lineage to the eukaryotic ancestor, but even more unexpectedly demonstrated that the descendants of the same archaeal lineage are still with us today (1-4). While Thor- and Lokiarchaeia have anaerobic lifestyles, Heimdallarchaeia appear capable of facultative aerobic metabolism, also possessing at least three types of light-activated rhodopsins (5). Despite the overall growing appreciation of these remarkable microbes, a pressing concern is that not a single member of Asgardarchaeota has yet been seen. We chose to tackle this issue by using CARD-FISH and epifluorescence microscopy techniques that have been previously successfully used for visualization of a wide array of microbes (6-9).

In order to design specific probes for a morphological characterization of Asgardarchaeota, we manually optimized the alignment of all 16S rRNA gene sequences classified as Asgardarchaeota in ARB (10) using the latest SILVA database (SSURef_NR99_132) (11) and constructed a RAxML tree (GTR-GAMMA model, 100 bootstraps (12)) for all high-quality near full-length sequences (Supplementary Figure 1). All attempts to construct a general probe targeting all Asgardarchaeota sequences (n=935) failed, likely due to the high diversity of this super-phylum. The impossibility to design such broad-range probes is not surprising, as similar difficulties have been reported for other diverse prokaryotic groups (e.g. Proteobacteria (8)). Even the widely used ‘general’ archaeal probe ARCH915 (13) is unspecific in this regard, overlapping only partially with Asgardarchaeota (80% coverage). This ‘general’ archaeal probe covers 85% of all archaea, 88.8% Lokiarchaea, but fails to detect most Heimdallarchaeia (only 13% targeted) and is thus not sufficiently reliable to detect Asgardarchaeota. Consequently, we decided to design specific probes for Heimdall- and Lokiarchaeia. Two probes and a set of competitors (14) were constructed targeting 93.4% of sequences affiliated to lineage loki1, the larger of the two specific branches of Lokiarchaeia, and 97% of Heimdallarchaeia (Supplementary Figure 1; Supplementary Tables 1, 2; for details see Supplementary methods).

We tested these probes in sediment samples from two sites from where recently several Asgardarchaeota genomes were recovered by metagenomics (Lakes Tekirghiol and Amara, South Eastern Romania) (5). Seven sediment layers (0-70 cm, in 10-cm ranges) were recovered from Lake Tekirghiol, and the top 10 cm was sampled in Lake Amara. Samples were fixed with formaldehyde, treated by sonication, vortexing and centrifugation to detach cells from sediment particles (15) and aliquots were filtered onto white polycarbonate filters (0.2 µm pore size, Millipore). CARD-FISH was conducted as previously described with fluorescein labelled tyramides (15). Filters were counterstained with DAPI and inspected by epifluorescence microscopy (for details see Supplementary methods).

Both phyla were rare and appeared completely absent below depths of 40 cm. All observed Heimdallarchaeia were similar in cell size (2.0±0.5 µm length × 1.4±0.4 µm width, n=23) and of conspicuous shape with DNA condensed (0.8±0.2 × 0.5±0.2 µm) at the center of the cells (Figure 1, Supplementary Figures 2, 4, 6), which is rather atypical for prokaryotes. In contrast, Lokiarchaeia presented diverse shapes and sizes and we could distinguish at least three distinct morphotypes. The most common Lokiarchaeia were small-medium sized, ovoid cells (2.0±0.5 × 1.4±0.3 µm, n=30, Figure 2 a-c, Supplementary Figure 3, 4, 5) that were found in different depth layers in Lake Tekirghiol (0-10 cm, 10-20 cm, 20-30 cm) and in the top 10 cm sample from Lake Amara. A single large round cell (3.8 × 3.6 µm, Figure 2 d-f) with bright fluorescence signal and condensed DNA at the center was detected in Lake Amara, and large rods/filaments (12.0±4.3 × 1.4±0.5 µm, n=6, Figure 2 g-i, Supplementary Figures 3, 4, 5) with filamentous, condensed DNA (10.2±4.8 × 0.6±0.1 µm) were present at 30-40 cm sediment depth in Lake Tekirghiol and in 0-10 cm depth in Lake Amara. The variety of Lokiarchaeia morphologies most likely reflects the higher sampling of the phylogenetic diversity within this phylum. Precise quantification was hampered by the very low abundances in the analyzed samples, which corresponds well to low recoveries of 16S rRNA reads from metagenomes (Supplementary Figure 6)(4). Consequently, it is difficult to draw firm conclusions on sediment depth preferences or rule out additional morphotypes of Lokiarchaeia.

**Figure 1:**
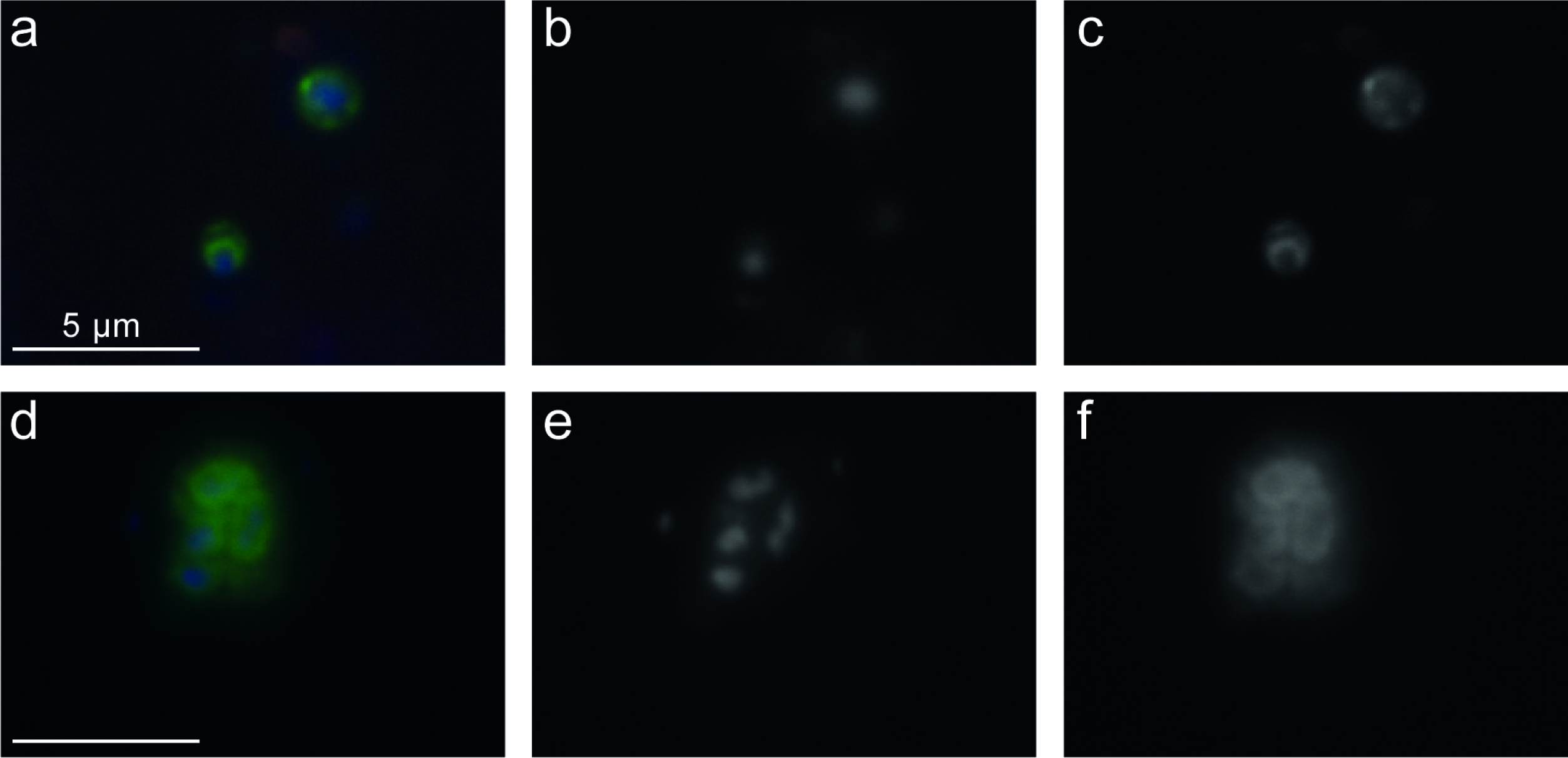
CARD-FISH imaging of Heimdallarchaeia hybridized with probe heim-526. The left panels (a, d) display overlay images of probe signal (green), DAPI staining (blue) and autofluorescence (red), the middle panels (b, e) DAPI staining of DNA, the right panels (c, f) CARD-FISH staining of proteins. Individual microphotographs of autofluorescent objects are not displayed because of low intensities and no interference with probe signals (see Figure S4). The scale bar (5 μm) in the left images applies to all microphotographs. Additional images of Heimdallarchaeia can be found in Supplementary Figure S2.

**Figure 2:**
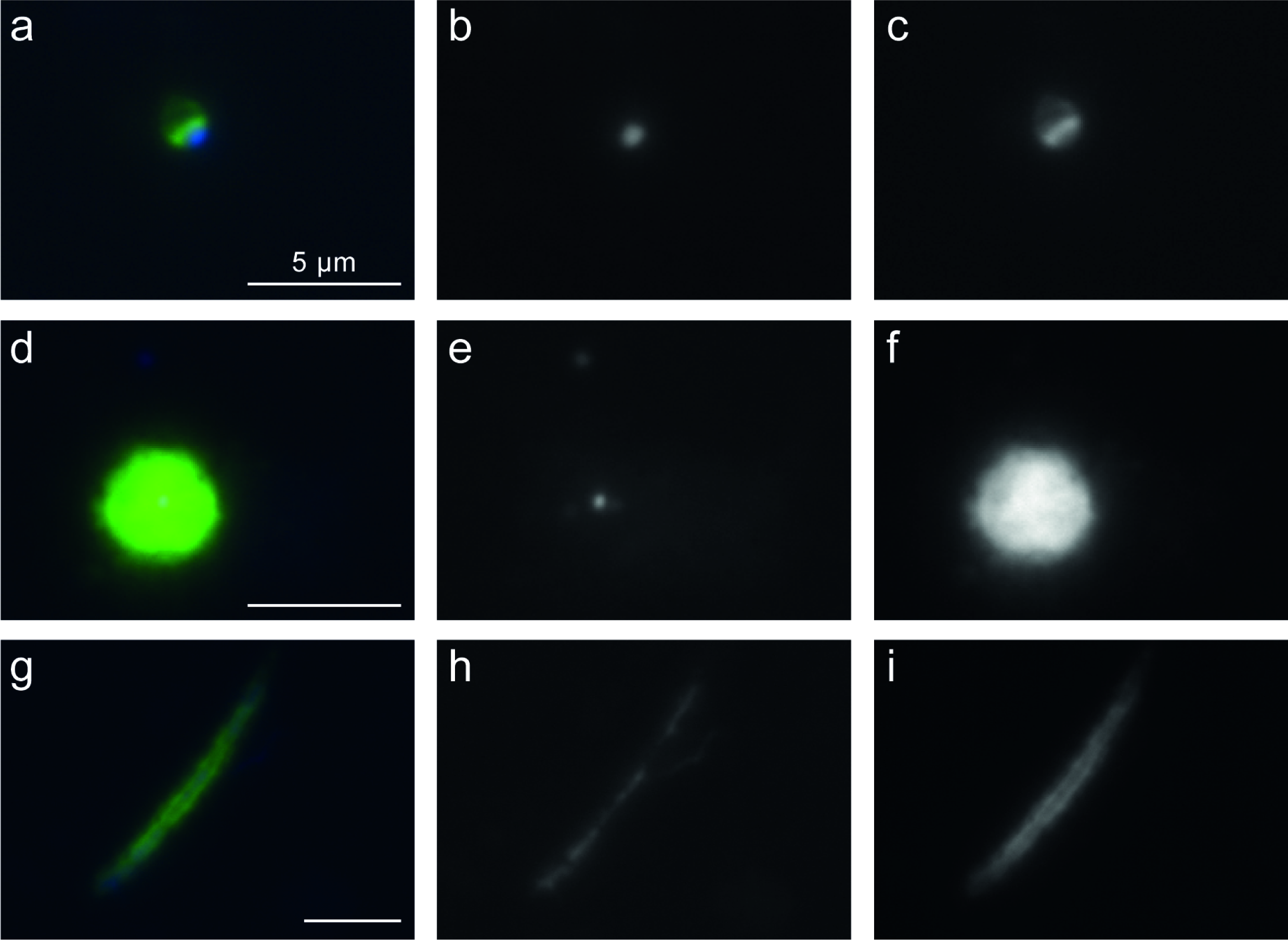
CARD-FISH imaging of Lokiarchaeia hybridized with probe loki1-1184. Three different morphotypes are displayed: (a-d) small-medium sized ovoid cells, (d-f) large round cell, and (g-i) large filamentous cells. The left panels (a, d, g) display overlay images of probe signal (green), DAPI staining (blue) and autofluorescence (red), the middle panels (b, e, h) DAPI staining of DNA, the right panels (c, f, i) CARD-FISH staining of proteins. Individual microphotographs of autofluorescent objects are not displayed because of low intensities and no interference with probe signals (see Figure S4). The scale bar (5 μm) in the left images applies to all microphotographs. Additional images of Lokiarchaeia can be found in Supplementary Figure S3.

During microscopic inspections, we carefully checked for potential non-specific or autofluorescent signals at wavelengths not interfering with the probe signal and found no overlap for any of the inspected cells. A set of negative controls were conducted to rule out false-positive signals due to unspecific binding of dye or nucleic acid components of probes by using a nonspecific probe (NON338, Supplementary Figure 7)(16). To avoid false-positive signals from cellular peroxidases, we performed additional control experiments including the CARD reaction only (without probes). All these control treatments resulted in low, unspecific background signals (comparable to those obtained in samples with probes for Asgardarchaeota), but no obvious staining of cells (Supplementary Figures 4, 7). Further evidence of specificity was seen in all cells hybridized with the Heimdallarchaeia probe, both the shapes and staining patterns coupled to DAPI were remarkably consistent.

While tempting, in absence of strong supporting evidence it would be too premature to conclude whether the condensed DNA, particularly in Heimdallarchaeia cells is indicative of a proto-nucleus. Microscopic images of bacterial cells with apparently eukaryotic-features have been misinterpreted before, e.g. in the case of the phylum Planctomycetes (17).

Examining these first images of Asgardarchaeota, we could not avoid to note the unusual happenstance of their naming with shapes and ecology. Their initial baptism after mythological characters from Norse mythology (Odin, Thor, Loki and Heimdall (3, 4)) has been unusually prescient (at least for Loki- and Heimdallarchaeia). The many-faced Loki lives up to the name under microscopic scrutiny (Figure 2, Supplementary Figure 3), and Heimdallarchaeia (Heimdall, ‘the one who illuminates the world’), appear to be if not light-bringer, at least light-sensitive (5). The unexpected parallels extend even further, Heimdall as gatekeeper between the realms of Midgard and Asgard, reprises the role in a phylogenetic reincarnation as a transition of the archaeal proto-eukaryotes from the dark, anoxic domain of Loki to the sun-lit, oxic world of present day eukaryotes (5).

## Acknowledgements

M.M.S was supported by the research grant 19-23469S (Grant Agency of the Czech Republic). A-Ş.A. was supported by the research grants 17-04828S (Grant Agency of the Czech Republic) and MSM200961801 (Academy of Sciences of the Czech Republic). P-A.B and Z.G.K were supported by the research grant PN-III-P4-ID-PCE-2016-0303 (Romanian National Authority for Scientific Research). H.B.L. was supported by the research grants PN-III-P4-ID-PCE-2016-0303 (Romanian National Authority for Scientific Research) and STAR-UBB Advanced Fellowship-Intern (Babeş-Bolyai University). R.G. was supported by the research grant 17-04828S (Grant Agency of the Czech Republic).

## Competing Interests

The authors declare no competing financial interests in relation to the work described.

## Supplementary information

### Supplementary methods

During *in silico* testing of the designed probes for specificity and group coverage, we identified two sequences that behaved aberrantly, i.e. could theoretically be regarded as outgroup hits for probe loki1-1184 (GU363076 and EU731577, **Supplementary Table 2**). However, these were discarded after closer examination because of pintail values of 20 and 0, respectively, indicating a chimeric origin (1). Pintail values are measures of chimeric nature for rRNA sequences in the databases, a value closer or equal to 100 indicating a non-chimeric sequence (1). Additional outgroup candidates for probe loki1-1184 (KU351219, AY133348, JQ817340, **Supplementary Table 2**) were initially located within Heimdallarchaeia in the basic tree provided by SILVA, but turned out to belong to Lokiarchaeia after alignment optimizations and RAxML tree reconstruction.

To be absolutely certain to avoid false-positive signals, i.e., to target no other organisms, we designed a set of competitor oligonucleotides that bind specifically to those rRNA sequences that have a single mismatch with our probes (2). We designed three distinct competitor probes for Heimdallarchaeota and two for Lokiarchaeota (**Supplementary Table 1**). Each competitor was used in the same concentrations as the CARD-FISH probes in order to prevent non-specific binding. The usage of specific probes together with competitors has been previously shown to work very well for visualizing cell-morphology and enumeration and was applied numerous times (3-7). Probes were tested *in silico* (8) and in the laboratory with different formamide concentrations in the hybridization buffer until stringent conditions were achieved (**Supplementary Table 1**).

We tested these probes in sediment samples from two sites from where recently several Asgardarchaeota genomes were recovered by metagenomics (Lakes Tekirghiol and Amara, Romania) (9). Sediment sampling was performed using a custom mud corer on 22 April 2018 at 12:00 in Tekirghiol Lake, Romania, (44°03.19017 N, 28°36.19083 E) and Amara Lake (44°36’N, 27°20’E, 23 April 2018, 14:00). Seven sediment layers (0-70 cm, in 10-cm ranges) were sampled in Lake Tekirghiol, and the top 10 cm was sampled in Lake Amara. Samples were fixed with formaldehyde for 1 h and washed three times with 1 × PBS, with centrifugation at 16000 g for 5 min between washes, and a final resuspension in a 1:1 mixture of PBS and ethanol. A treatment of sonication (20 sec, minimum power) on ice, vortexing and centrifugation to detach cells from sediment particles was applied (10) and aliquots diluted with PBS were filtered onto white polycarbonate filters (0.2 µm pore size, Millipore). CARD-FISH was conducted as previously described with fluorescein labelled tyramides and included permeabilization steps with lysozyme and achromopeptidase and an inactivation step for cellular peroxidases with methanol (10). Filters were counterstained with DAPI and inspected by epifluorescence microscopy (Zeiss Imager.M1) with filter sets for DAPI (filter set 01: BP 365/12, FT 395, LP 397), fluorescein (filter set 10: BP 450-490, FT 510, BP 515-565), and autofluorescence (filter set 15: BP 546/12, FT 580, LP 590). Micrographs of CARD-FISH stained cells were recorded with a highly sensitive charge-coupled device (CCD) camera (Vosskühler) at fixed exposure times (70 and 100 ms for DAPI, 100 and 200 ms for CARD-FISH and 100 and 400 ms for autofluorescence for magnifications of 400 × and 1000 × respectively) and cell sizes were estimated with the software LUCIA (Laboratory Imaging Prague, Czech Republic).

---

**Supplementary Figure 1:**
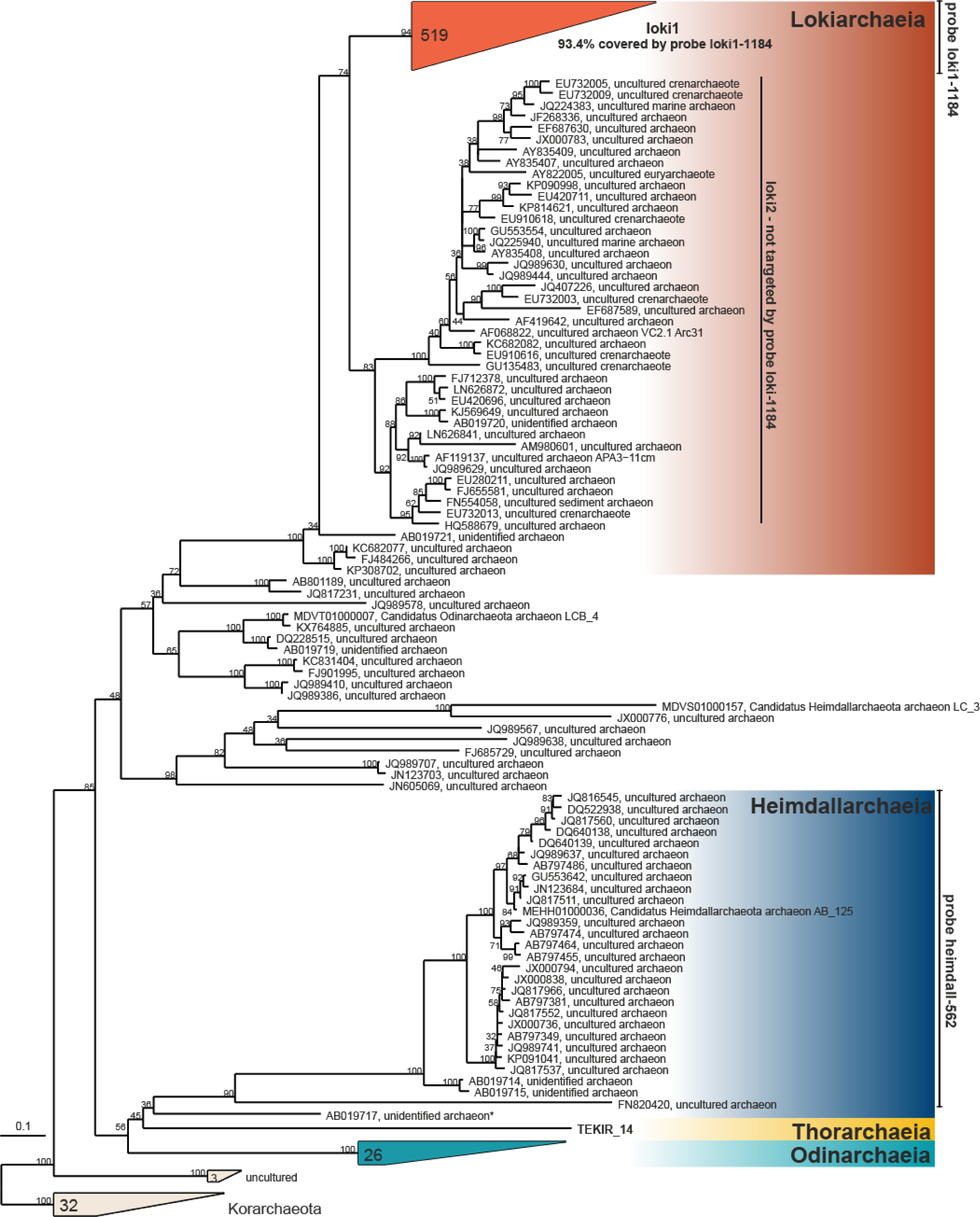
RAxML tree (GTR-GAMMA model, 100 bootstraps) of 16S rRNA genes of Asgardarchaeota with target hits of probes loki1-1184 and heim-562. Branches with bootstrap supports <30% were collated to multifurcations. The number of sequences is given for collapsed branches; an asterisk indicates the sequence not targeted by probe heim-562.

**Supplementary Figure 2:**
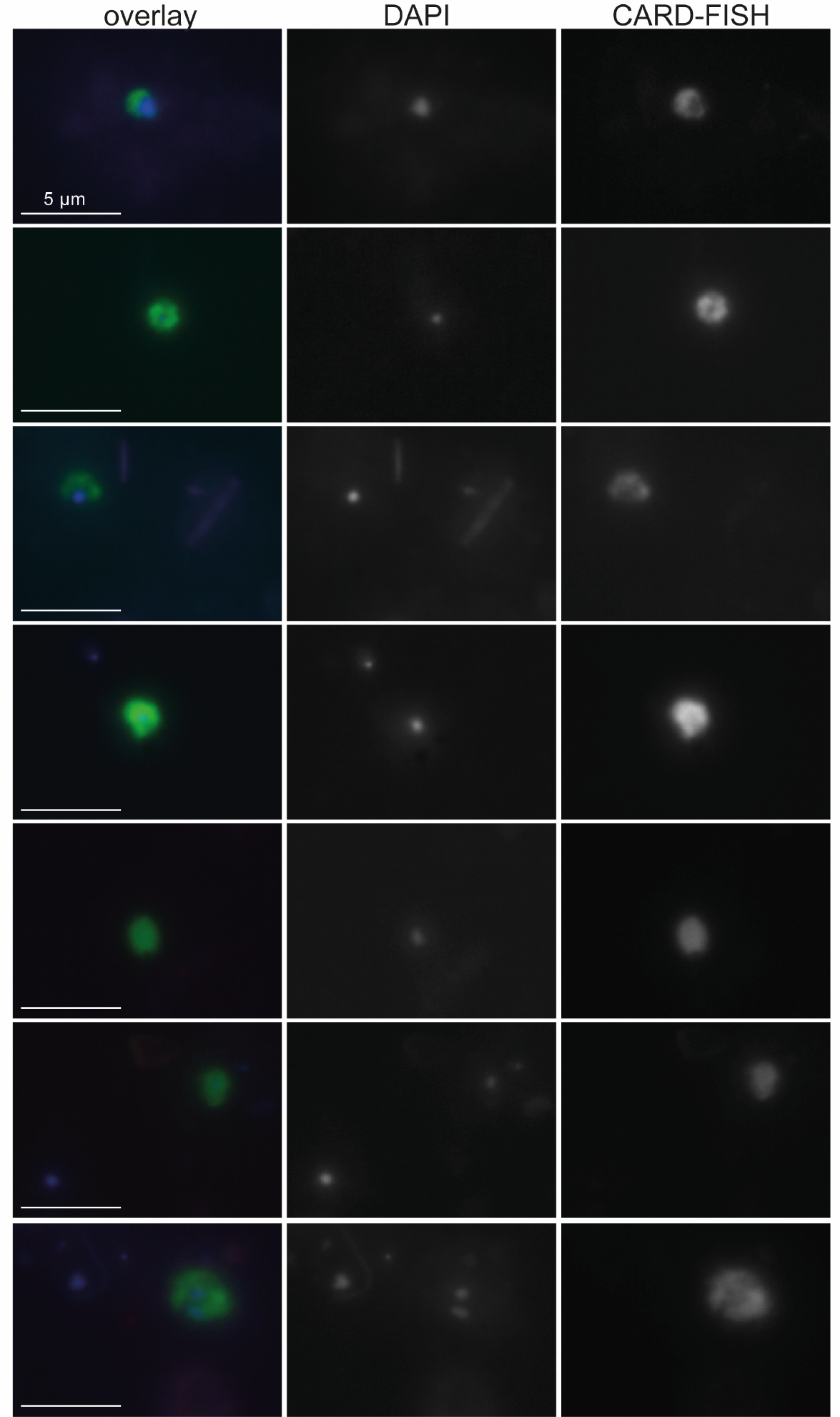
CARD-FISH images of Heimdallarchaeia (probe heim-526). The left panels display overlay images of probe signal (green), DAPI staining (blue) and autofluorescence (red), the middle panels DAPI staining of DNA, the right panels CARD-FISH staining of proteins. Individual microphotographs of autofluorescent objects are not displayed because of low intensities and no interference with probe signals.

**Supplementary Figure 3.**
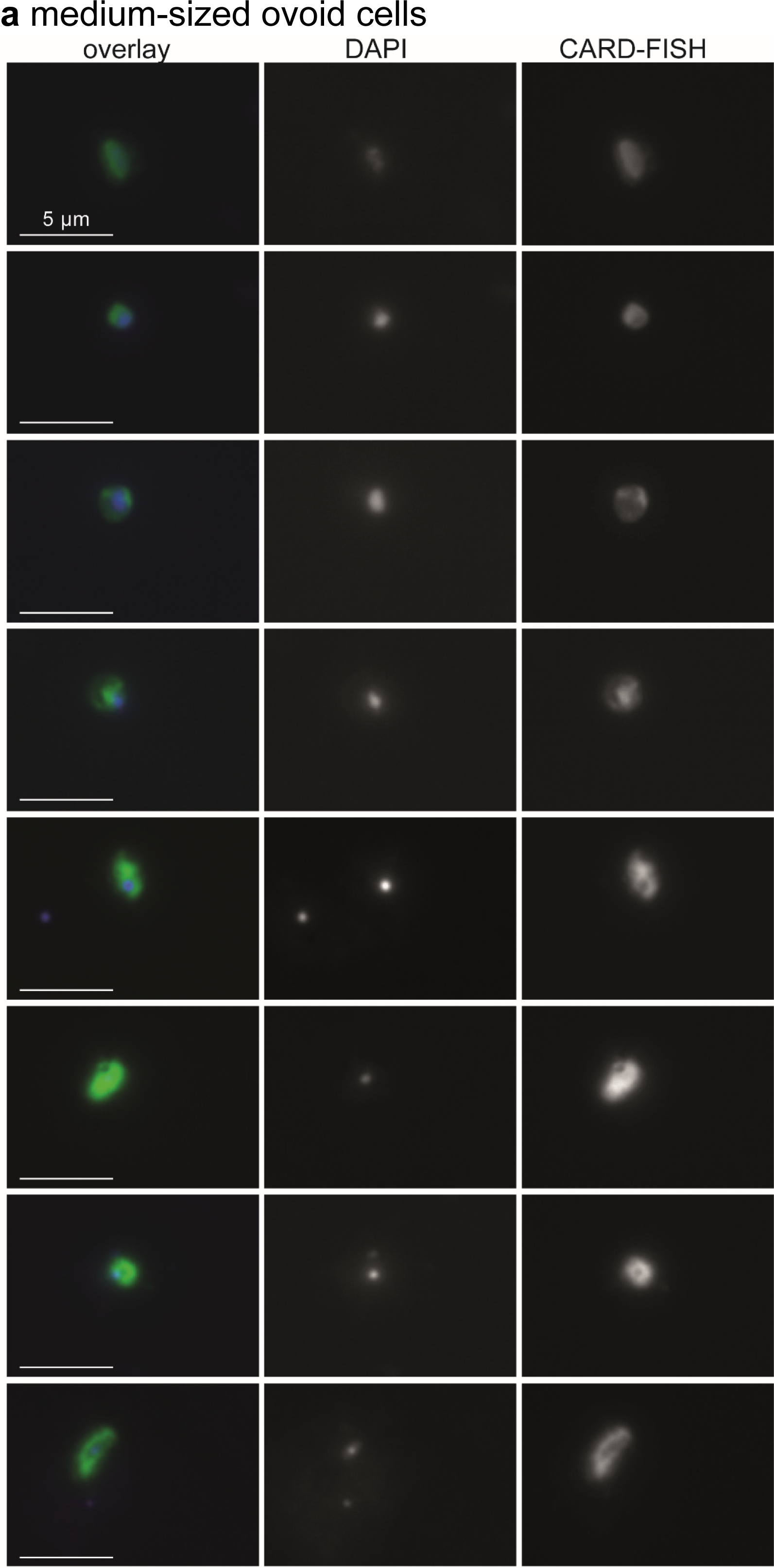
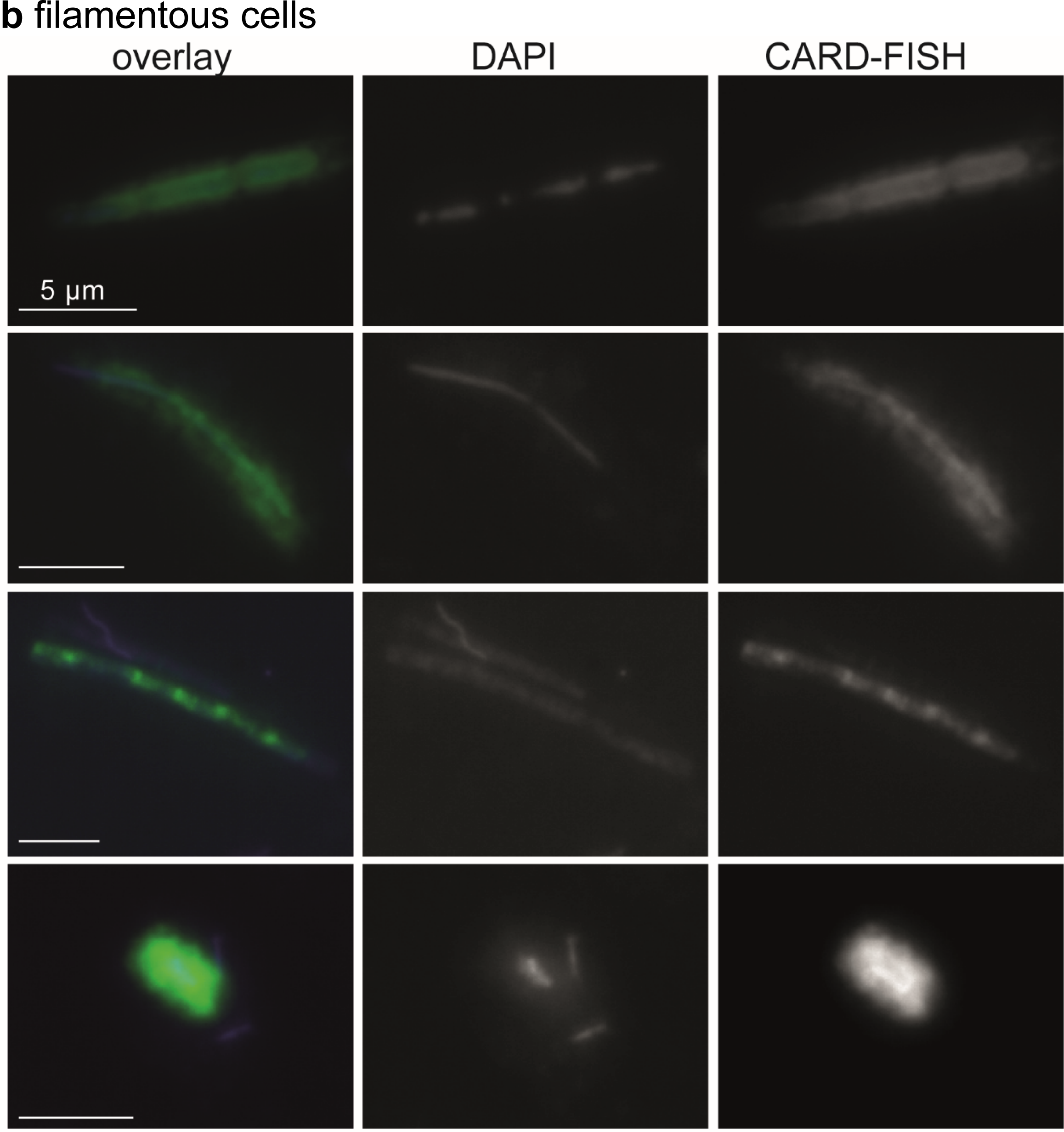
CARD-FISH imaging of two distinct morphotypes of Lokiarchaeia (probe loki1-1184), (a) medium-sized ovoid cells and (b) filamentous cells. The left panels display overlay images of probe signal (green), DAPI staining (blue) and autofluorescence (red), the middle panels DAPI staining of DNA, the right panels CARD-FISH staining of proteins. Individual microphotographs of autofluorescent objects are not displayed because of low intensities and no interference with probe signals. The scale bar (5 μm) in the left images applies to all microphotographs.

**Supplementary Figure 4:**
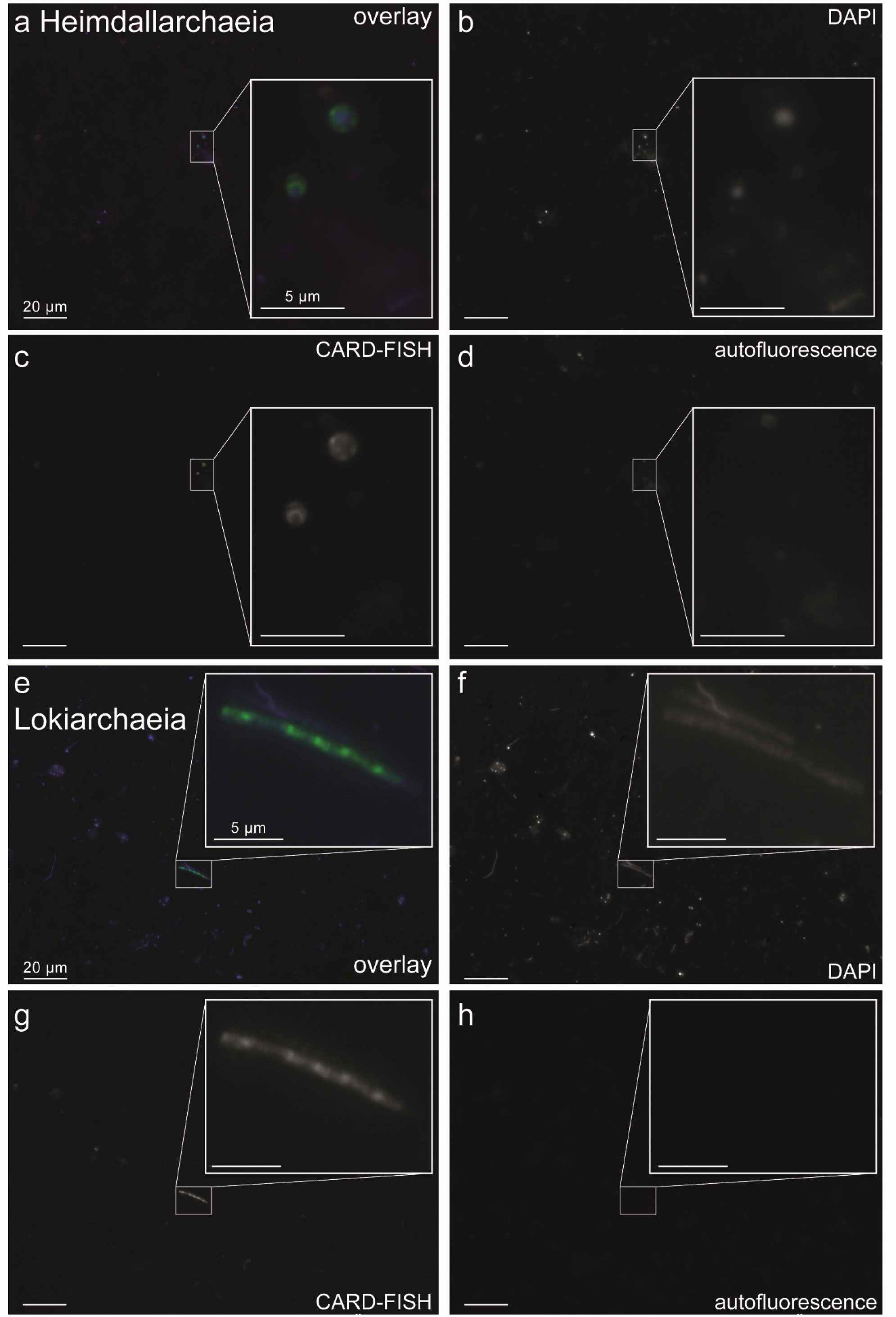
Images of Heimdallarchaeia (a-d) and Lokiarchaeia (e-h) at 400 × magnification with inserts taken at 1000 × magnification. (a, e): Overlay images of probe signal (green), DAPI staining (blue) and autofluorescence (red); (b, f): DAPI staining; (c, g): CARD-FISH staining; (d, h): autofluorescence. Images were recorded at fixed exposure times (70 or 100 ms for DAPI, 100 or 200 ms for CARD-FISH and 100 or 400 ms for autofluorescence for 400 × or 1000 × magnification, respectively).

**Supplementary Figure 5:**
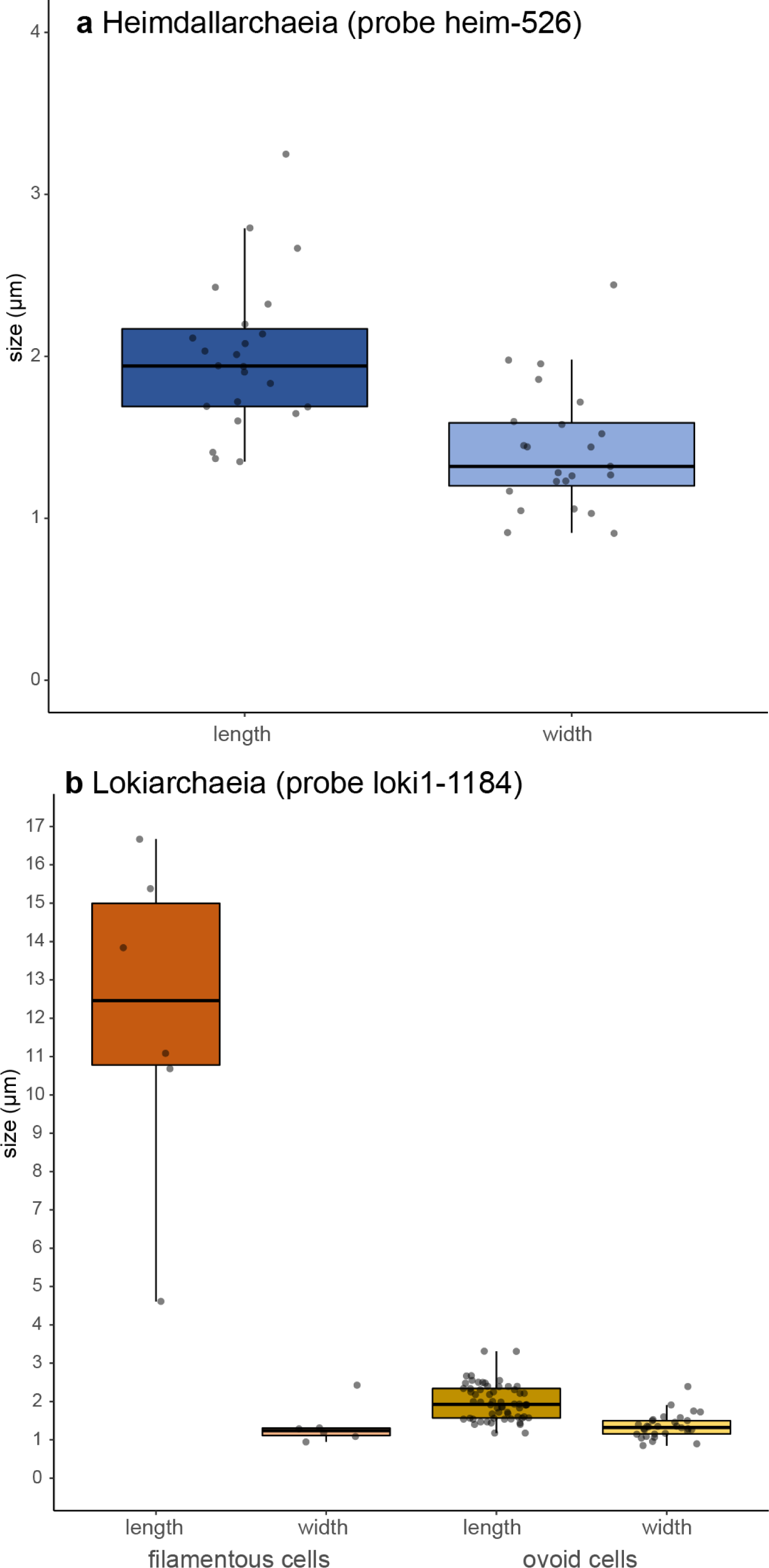
Cell sizes (lengths and widths) of (a) Heimdallarchaeia (n=23) and (b) two different morphotypes of Lokiarchaeia (filaments, n=6; small-medium sized ovoid cells, n=30).

**Supplementary Figure 6:**
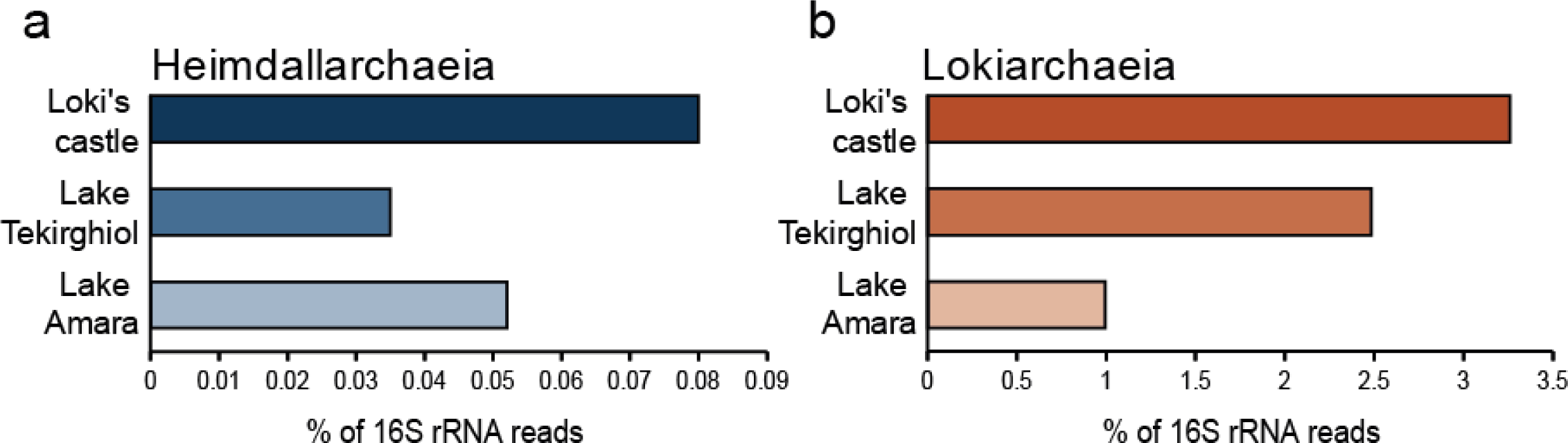
Abundance estimation of recovered 16S rRNA reads affiliated with Heimdallarchaeia (a) and Lokiarchaeia (b) from metagenomes from Loki’s castle, Lake Tekirghol and Lake Amara. Shotgun metagenomes from Amara (SRA7615342) and Tekirghiol (SRA7614767) lakes, as well as the published SRX684858 from Loki’s castle were subsampled to 20 million sequences. Each subset was queried for putative RNA sequences against the non-redundant SILVA SSURef_NR99_132 database, that was clustered at 85% sequence identity. Identified putative 16S rRNA sequences (e-value < 1e-5) were screened using SSU-ALIGN. Resulting bona fide 16S rRNA sequences were compared by blastn (e-value <1e-5) against the curated SILVA SSURef_NR99_132 database. Matches with identity ≥ 80% and alignment length ≥90 bp were considered for downstream analyses. Sequences assigned to Loki- and Heimdallarchaeia were used to calculate abundances for these taxa in their originating environments.

**Supplementary Figure 7:**
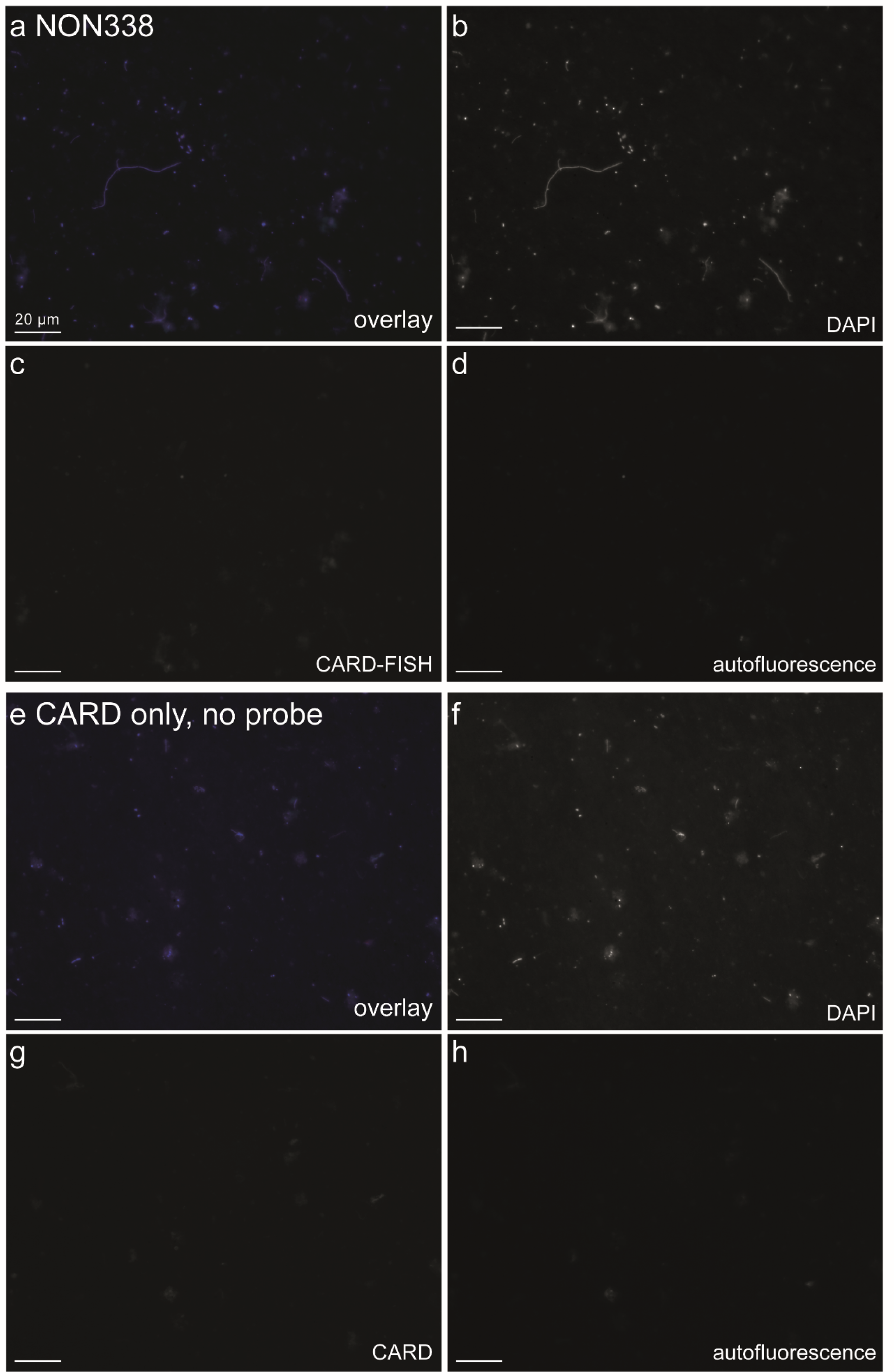
Control treatments with (a-d) nonsense probe NON338 and (e-h) the CARD reaction only without using a probe. Overlay images of probe signal (green), DAPI staining (blue) and autofluorescence (red); (b, f): DAPI staining; (c, g): CARD-FISH or CARD only staining; (d, h): autofluorescence. Images were recorded at fixed exposure times (70 ms for DAPI, 100 ms for CARD-FISH and 100 ms for autofluorescence).

## References

1. Cox CJ, Foster PG, Hirt RP, Harris SR, Embley TM. The archaebacterial origin of eukaryotes. Proceedings of the National Academy of Sciences. 2008;105(51):20356–61.

2. Lake JA, Henderson E, Oakes M, Clark MW. Eocytes: a new ribosome structure indicates a kingdom with a close relationship to eukaryotes. Proceedings of the National Academy of Sciences. 1984;81(12):3786–90.

3. Spang A, Saw JH, Jørgensen SL, Zaremba-Niedzwiedzka K, Martijn J, Lind AE, et al. Complex archaea that bridge the gap between prokaryotes and eukaryotes. Nature. 2015;521:173.

4. Zaremba-Niedzwiedzka K, Caceres EF, Saw JH, Bäckström D, Juzokaite L, Vancaester E, et al. Asgard archaea illuminate the origin of eukaryotic cellular complexity. Nature. 2017;541:353.

5. Bulzu P-A, Andrei A-S, Salcher MM, Mehrshad M, Inoue K, Kandori H, et al. Casting light on Asgardarchaeota metabolism in a sunlit microoxic niche. Nat Microbiol. accepted.

6. Andrei A-S, Salcher MM, Mehrshad M, Rychtecký P, Znachor P, Ghai R. Niche-directed evolution modulates genome architecture in freshwater Planctomycetes. The ISME Journal. 2019.

7. Neuenschwander SM, Ghai R, Pernthaler J, Salcher MM. Microdiversification in genome-streamlined ubiquitous freshwater Actinobacteria. ISME J. 2018;12:185.

8. Amann R, Fuchs BM. Single-cell identification in microbial communities by improved fluorescence in situ hybridization techniques. Nat Rev Microbiol. 2008;6(5):339–48.

9. Mußmann M, Brito I, Pitcher A, Sinninghe Damsté JS, Hatzenpichler R, Richter A, et al. Thaumarchaeotes abundant in refinery nitrifying sludges express amoA but are not obligate autotrophic ammonia oxidizers. Proc Natl Acad Sci USA. 2011;108(40):16771–6.

10. Ludwig W, Strunk O, Westram R, Richter L, Meier H Yadhukumar, et al. ARB: a software environment for sequence data. Nucl Acid Res. 2004;32(4):1363–71.

11. Pruesse E, Quast C, Knittel K, Fuchs BM, Ludwig W, Peplies J, et al. SILVA: a comprehensive online resource for quality checked and aligned ribosomal RNA sequence data compatible with ARB. Nucl Acid Res. 2007;35(21):7188–96.

12. Stamatakis A, Ludwig T, Meier H. RAxML-II: a program for sequential, parallel and distributed inference of large phylogenetic. Concurr Comput-Pract Exp. 2005;17(14):1705–23.

13. Stahl DA, Amann R. Development and application of nucleic acid probes. In: Stackebrandt E, Goodfellow M, editors. Nucleic acid techniques in bacterial systematics. Chichester, England: John Wiley & Sons Ltd.; 1991. p. 205–48.

14. Fuchs B, Glöckner F, Wulf J, Amann R. Unlabeled helper oligonucleotides increase the in situ accessibility to 16S rRNA of fluorescently labeled oligonucleotide probes. Appl Environ Microbiol. 2000;66:3603–7.

15. Ishii K, Mußmann M, MacGregor BJ, Amann R. An improved fluorescence in situ hybridization protocol for the identification of bacteria and archaea in marine sediments. FEMS Microbiol Ecol. 2004;50(3):203–13.

16. Wallner G, Amann R, Beisker W. Optimizing fluorescent in situ hybridization with rRNA-targeted oligonucleotide probes for flow cytometric identification of microorganisms. Cytometry. 1993;14(2):136–43.

17. Fuerst JA, Sagulenko E. Beyond the bacterium: planctomycetes challenge our concepts of microbial structure and function. Nat Rev Microbiol. 2011;9:403.

## Additional references

1. Ashelford K, Chuzhanova N, Fry J, Jones A, Weightman A. At least 1 in 20 rRNA sequence record currently held in public repositories is estimated to contain substantial anomalies. Appl Environ Microbiol. 2005;71:7724–36.

2. Fuchs B, Glöckner F, Wulf J, Amann R. Unlabeled helper oligonucleotides increase the in situ accessibility to 16S rRNA of fluorescently labeled oligonucleotide probes. Appl Environ Microbiol. 2000;66:3603–7.

3. Teeling H, Fuchs BM, Becher D, Klockow C, Gardebrecht A, Bennke CM, et al. Substrate-controlled succession of marine bacterioplankton populations induced by a phytoplankton bloom. Science. 2012;336(6081):608–11.

4. Mußmann M, Brito I, Pitcher A, Sinninghe Damsté JS, Hatzenpichler R, Richter A, et al. Thaumarchaeotes abundant in refinery nitrifying sludges express amoA but are not obligate autotrophic ammonia oxidizers. Proc Natl Acad Sci USA. 2011;108(40):16771–6.

5. Andrei A-Ş, Salcher MM, Mehrshad M, Rychtecký P, Znachor P, Ghai R. Niche-directed evolution modulates genome architecture in freshwater Planctomycetes. The ISME Journal. 2019.

6. Neuenschwander SM, Ghai R, Pernthaler J, Salcher MM. Microdiversification in genome-streamlined ubiquitous freshwater Actinobacteria. ISME J. 2018;12:185.

7. Salcher MM, Posch T, Pernthaler J. *In situ* substrate preferences of abundant bacterioplankton populations in a prealpine freshwater lake. ISME J. 2013;7:896–907.

8. Yilmaz LS, Parnerkar S, Noguera DR. mathFISH, a web tool that uses thermodynamics-based mathematical models for in silico evaluation of oligonucleotide probes for fluorescence *in situ* hybridization. Appl Environ Microbiol. 2011;77(3):1118–22.

9. Bulzu P-A, Andrei A-S, Salcher MM, Mehrshad M, Inoue K, Kandori H, et al. Casting light on Asgardarchaeota metabolism in a sunlit microoxic niche. Nat Microbiol. accepted.

10. Ishii K, Mußmann M, MacGregor BJ, Amann R. An improved fluorescence in situ hybridization protocol for the identification of bacteria and archaea in marine sediments. FEMS Microbiol Ecol. 2004;50(3):203–13.

